# Inside Information: Systematic within-node connectivity changes observed across tasks or groups

**DOI:** 10.1101/2021.07.07.451429

**Authors:** Wenjing Luo, R. Todd Constable

## Abstract

Mapping the human connectome and understanding its relationship to brain function holds tremendous clinical potential. The connectome has two fundamental components: the nodes and the connections between them. While much attention has been given to deriving atlases and measuring the connections between nodes, there have been no studies examining the networks within nodes. Here we demonstrate that each node contains significant connectivity information, that varies systematically across task-induced states and subjects, such that measures based on these variations can be used to classify tasks and identify subjects. The results are not specific for any particular atlas but hold across different atlas resolutions. To date, studies examining changes in connectivity have focused on edge changes and assumed there is no useful information within nodes. Our findings illustrate that for typical atlases, within-node changes can be significant and may account for a substantial fraction of the variance currently attributed to edge changes.

## Introduction

Since the concept of the human connectome was first introduced by Sporns in 2005 ^1^, there has been tremendous interest in mapping the connectome and understanding its relationship to brain function in both the healthy and diseased brain. The connectome, whether it is constructed based on structural or functional connections (or both), requires two primary ingredients: the nodes and the connections between them. Once the nodes are defined, the connectome is constructed, by measuring the connections between all pairs of nodes. The connectome is a connectivity matrix, with both the rows and the columns representing the nodes, and the cells representing the strength of connection (called edges) between pairs of nodes. Magnetic resonance imaging (MRI), generally does not have directional information, and so the matrix is diagonally symmetric. The node is a fundamental component of the connectome, and these are usually defined by applying a brain atlas and averaging the BOLD time-course signals within each node prior to calculating the connectivity. Here we focus on functional connectivity (in contrast to structural connectivity) defined as the Pearson’s r correlation of BOLD signals from pairs of nodes^2^. Once the matrix is obtained numerous approaches have been developed to measure changes in the connectome between groups and conditions, or in many cases graph theory approaches are applied to quantitatively summarize node and network properties ^3^. With such measures, connectivity changes can be quantified between brain states and/or groups or as a function of some behavioral or clinical measure. Previous studies have shown correlations between functional connectivity and subject traits, such as sex ^4, 5^, age ^5, 6^, fluid intelligence ^7, 8^ and attention ^9, 10, 11^. Network measures have also been related to subject features ^12, 13, 14^ and brain states ^15, 16^.

While there is consensus on, and a large literature supporting, the application of graph theory to neuroimaging data, it is generally ill-suited to such analyses. First there is no consensus on what to use for a brain atlas to define nodes. Secondly, unlike say disease propagation or the tracking of friends in social networks, whether or not two nodes are connected (the disease was transmitted between individuals, or two people are or are not friends on Facebook, for example) neuroimaging data typically does not provide a binary framework easily defining whether or not two nodes are connected. Thus, data must be binarized with an arbitrarily threshold to define a graph. Here we focus on subtleties associated with the application of an atlas to define the nodes.

Parcellation of the cortex and subcortex is not only a key first step in defining nodes for network analysis. Because of the central importance of such atlases, brain parcellation has been an active area of research and a wide range of approaches and modalities have been used to attempt to define the ideal atlas (for a review see ^17^). While early atlases were based on cytoarchitecture (ala Brodmann ^18^) with the advent of 3D imaging approaches, atlases have been built based on morphometric information, white matter microstructure obtained through diffusion tensor imaging (DTI) MRI, myelography, task-based functional MRI, functional connectivity, and gene expression data. Recent work has yielded atlases based on individuals ^19, 20^, groups of subjects ^21, 22, 23, 24, 25, 26^, based on anatomical MRI data ^27, 28^, functional data ^23, 24, 25, 26^, or a combination of both ^21, 22^. These atlases vary in their resolution (number of nodes in each atlas), coverage, and spatial perspective (volume-based or surface-based). The choice of atlas is somewhat critical as it has been shown that the results of functional connectivity analyses are atlas-dependent ^29, 30^ as are the results of network analysis ^30, 31, 32^ when using functional connectivity data.

In addition to the lack of consensus on which atlas to use, recent work^33^ has highlighted how node boundaries defined by functional connectivity are flexible within a fixed anatomic framework. While it has long been known that there is sub-specialization in cortical regions, this raises two fundamental questions with respect to connectivity analyses: 1) Is there an ideal atlas for defining functional nodes; 2) Does it make sense to combine structural data (which is fixed at the functional time scale) with functional data which is dynamic and flexible in its organization. Node reorganization over short temporal windows has been demonstrated in non-overlapping whole-brain parcellations ^34^. There is considerable support for this flexible organization in the literature from studies using a variety of methods ^35, 36, 37, 38, 39^. Recent functional connectivity work has shown that quantitative results in graph theory measures when comparing groups of subjects or comparing the same subjects across different task-conditions, change significantly if node reconfiguration is taken into account through flexible individualized- or group-brain-state-dependent parcellations ^40^. Yet fixed atlases are applied in most studies of the connectome, and the search continues for the ideal atlas ^41, 42, 43^.

Fundamentally, the use of an atlas comes with the implicit assumption that the node functions as a uniform entity. If this assumption is wrong and within-node connectivity does change, such changes can be incorrectly interpreted as edge changes. Node homogeneity, defined as the mean pair-wise correlation between voxel-level time series within the node, is in fact often used as a measure of goodness of parcellation with better parcellations having higher within node homogeneity. Maximal node homogeneity is achieved when the time-courses of the voxels within the node are all very similar, which would be the case with uniform neural activity throughout the node.

In this work we demonstrate that the neural activity is far from uniform in most nodes, and most importantly, that the homogeneity changes systematically with tasks. For a given atlas one can create a homogeneity vector of length equal to the number of nodes in the atlas. Using this vector we show, across a range of atlases, that there is considerable information in the vector and that there are reliable task-dependent changes in connectivity within the nodes defined by typical atlases. While previous work showed that with a flexible functional parcellation approach a node size vector could provide a measure that could be used to predict both the corresponding task during which the data were recorded as well as in-magnet task performance ^33^. Here we use node homogeneity to summarize between-voxel connectivity within-nodes and demonstrate that there is significant connectivity information contained within a node. The conclusion is that if one is interested in how functional connections in the brain change between tasks or groups, then all such changes (including within node changes) should be considered, and analyses should not be limited to only the possibility of edge changes.

## Results

In this work, homogeneity vectors containing the mean with-node connectivity for every node were obtained based on three fixed atlases at different resolution and used as features for task classification, sex classification, and subject identification. 493 subjects in the HCP dataset were used for task classification and subject identification. 422 subjects (211 males and 211 females) were used for sex classification. Homogeneity vectors based on individualized parcellations, state-specific parcellations and intersect parcellation were also used in the same classification and identification.

### 3.1 Task-induced brain states can be decoded from homogeneity vectors

Figure 1a demonstrates that the gradient boosting classification model can predict, at high accuracy, task labels of scans based on homogeneity vectors. The task classification accuracies are essentially the same for the three atlases applied. The accuracy obtained from the permutation test where we randomly assign task labels to scans is very low and consistent with the theoretical probability by chance (1/9).

**Figure 1.**
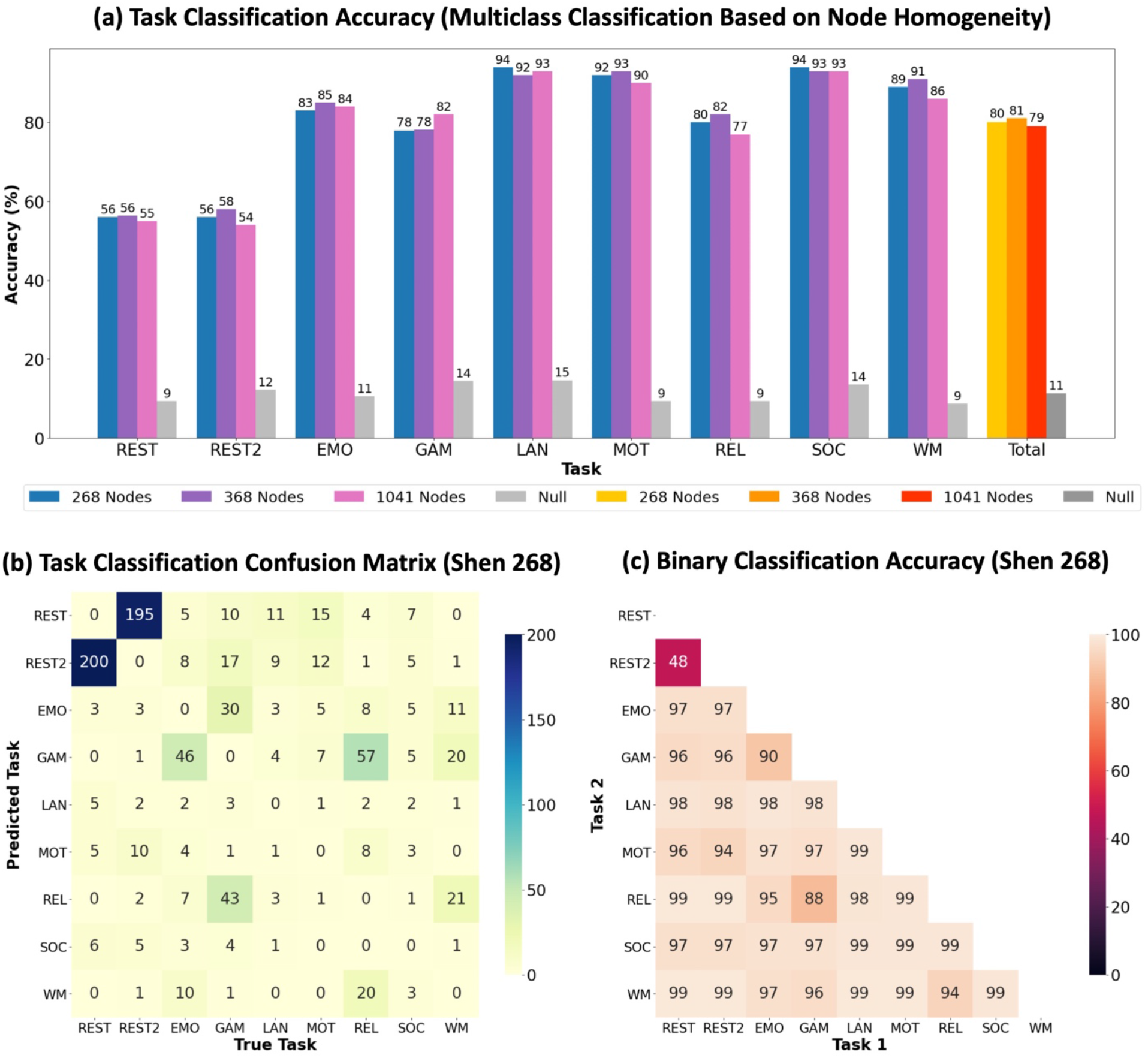
(a) The task classification accuracies for each task (the scans whose true labels are this particular task) and the total classification accuracy. The accuracies for three different parcellations, Shen 268, Shen 368, and Yeo 1041 are reported in different colors. The gray bars represent the accuracies obtained from the permutation test where the labels are randomly assigned to the scans. (b) The confusion matrix for task classification based on Shen 268. The numbers represent the number of scans whose true labels are shown on the x axis but are misclassified as the task on the y axis. Darker color means more misclassified scans. (c) The accuracies for binary task classification. Darker color means lower classification accuracy, i.e. more misclassified scans.

Figure 1b shows the confusion matrix for multiclass task classification with Shen 268, i.e. what the true labels and predicted labels are for the misclassified scans. The major confusion was between REST and REST2. Among a total of 494 REST scans, 200 were misclassified as REST2, and similarly, 195 REST2 scans were misclassified as REST. Other relatively frequently misclassified task pairs include GAMBLING and RELATIONAL, EMOTION and GAMBLING, RELATIONAL, and WM, in the order of decreasing frequency.

The accuracies of binary task classifications for all task pairs are presented in Figure 1c. The accuracy for REST vs. REST2 is 48% which is close to chance (50%). The accuracies for all the other task pairs are high and the lower ones among them are for GAMBLING vs. RELATIONAL, EMOTION vs. GAMBLING, RELATIONAL vs. WM, and REST2 vs. MOTOR. The task pairs with low binary classification accuracies are generally consistent with the pairs frequently get confused in multiclass classification.

### 3.2 Homogeneities of different nodes contribute unequally to task classification

To evaluate the importance of the homogeneity of each node, i.e. each element in the homogeneity vector for multiclass task classification, we used the impurity-based feature importance extracted from the classifiers. Figure 2a shows the importance ranking of each node’s homogeneity where the darker color represents higher importance. Figure 2b summarizes the lobe distribution of the twenty most important nodes for task classification. Most of the nodes are in the occipital lobe (45%) followed by the temporal lobe (25%) and the parietal lobe (20%). A few nodes are in the motor strip (10%).

**Figure 2.**
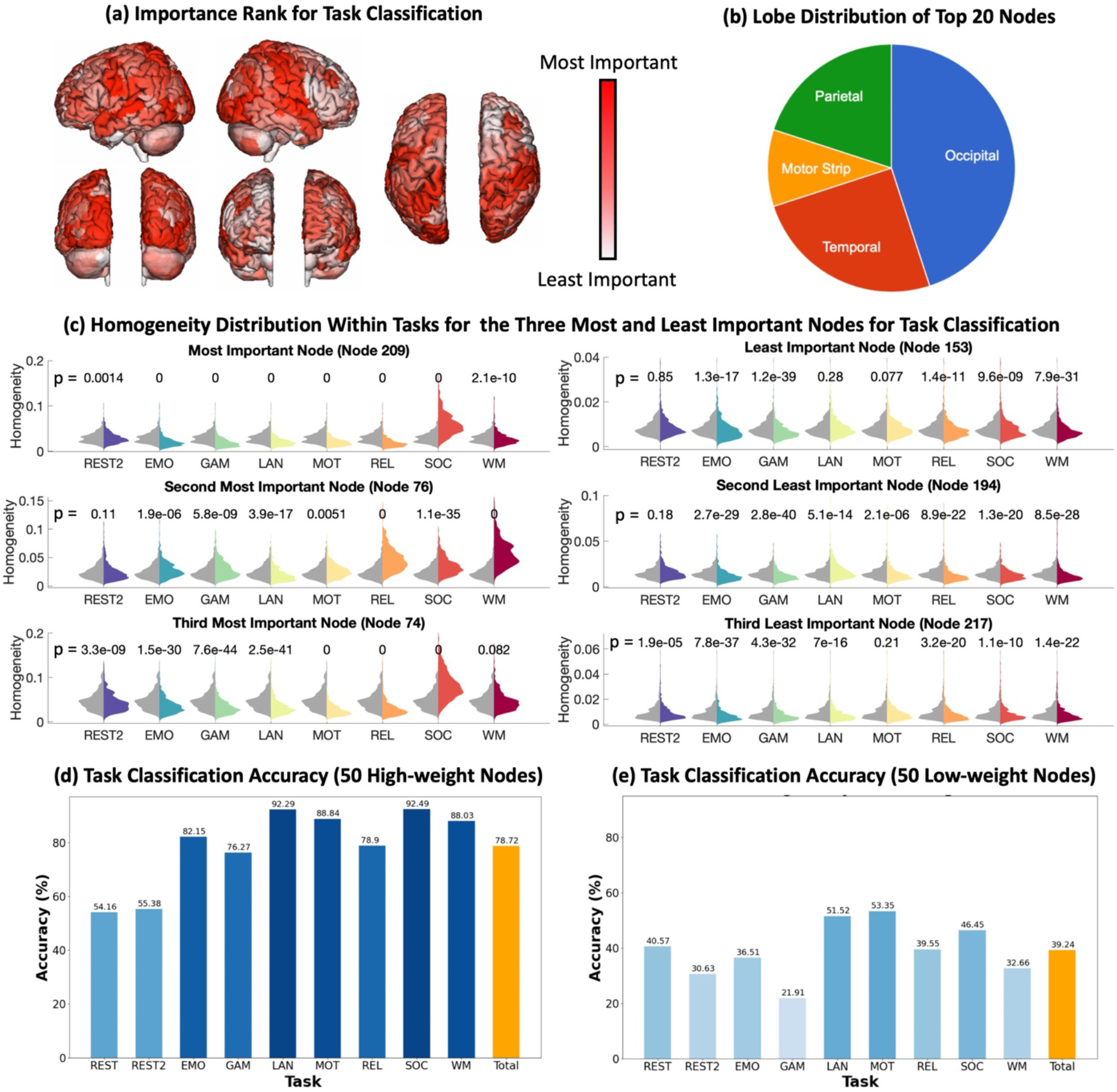
(a) The importance ranking of each node’s homogeneity for multiclass task classification. The darker color represents higher importance. (b) The lobe distribution of the twenty most important nodes for task classification. (c) The distributions of homogeneities for the three most important nodes and three least important nodes for multiclass task classification under different tasks. The gray distributions show the homogeneities under REST for 493 subjects and the distributions in other colors represent the homogeneities under other tasks labeled on the x axis. Paired t-test was used to compare the homogeneity distributions under REST and tasks. The p values are reported (above each pair of distribution). (d) The multiclass task classification accuracies for each task and the total accuracy using homogeneity vectors containing the fifty most node homogeneities. (e) The multiclass task classification accuracies for each task and the total accuracy using homogeneity vectors containing the fifty least node homogeneities.

We plotted the distributions of homogeneities for the three most important nodes and three least important nodes under different tasks to further explore how homogeneities vary with states. (Figure 2c) For the three most important nodes (left panel), the homogeneity distributions for REST (gray) are always significantly different from the distributions for other tasks at p<0.05, except for Node 76, REST vs. REST2. Some of the distributions for REST and tasks are especially visually distinguishable, such as Node 209, REST vs. SOCIAL, Node 76 REST vs. RELATIONAL and WM. For the three least important nodes, most of the distributions for REST and tasks are still significantly different, but more non-significant cases are observed and the p values are generally higher which means the significance levels are lower. It is also demonstrated in Figure 2c that the homogeneities for the three most important nodes are higher than the three least important nodes.

We also performed the same task classification described in 3.1 with homogeneities of a subset of the nodes, rather than all 268 nodes. Figure 2d shows the classification accuracies for each task and the total accuracy using homogeneity vectors containing the fifty most node homogeneities. The accuracies are comparable to the accuracies with complete homogeneity vectors reported in Figure 1a. However, with the fifty least important node homogeneities, the classification accuracies decrease by a lot as shown in Figure 2e. Although the accuracies are still above chance.

### 3.3 Between-voxel connectivity varies with brain states which leads to homogeneity changes

As shown in Figure 2c, the homogeneity of Node 209 is the most important feature for task classification and the homogeneity distributions are the most distinctive between REST and SOCIAL. To further investigate the between-voxel connectivity changes leading to such difference in homogeneity, we used one voxel in the center of Node 209 (marked in red in Figure 3) as the seed voxel and correlated its timeseries to the timeseries of all the other voxels in Node 209. The correlation coefficients (functional connectivity) are shown in Figure 3 for REST and SOCIAL. The result for a representative subject (subject 156536) is presented here. Only a small local region around the seed voxel has high connectivity with the seed for REST, partly because no spatial smoothing was applied. However, for SOCIAL, many more voxels within the node have high connectivity with the node voxel. Such an increase in between-voxel functional connectivity during SOCIAL can lead to the significantly higher homogeneity observed in Figure 2c.

**Figure 3.**
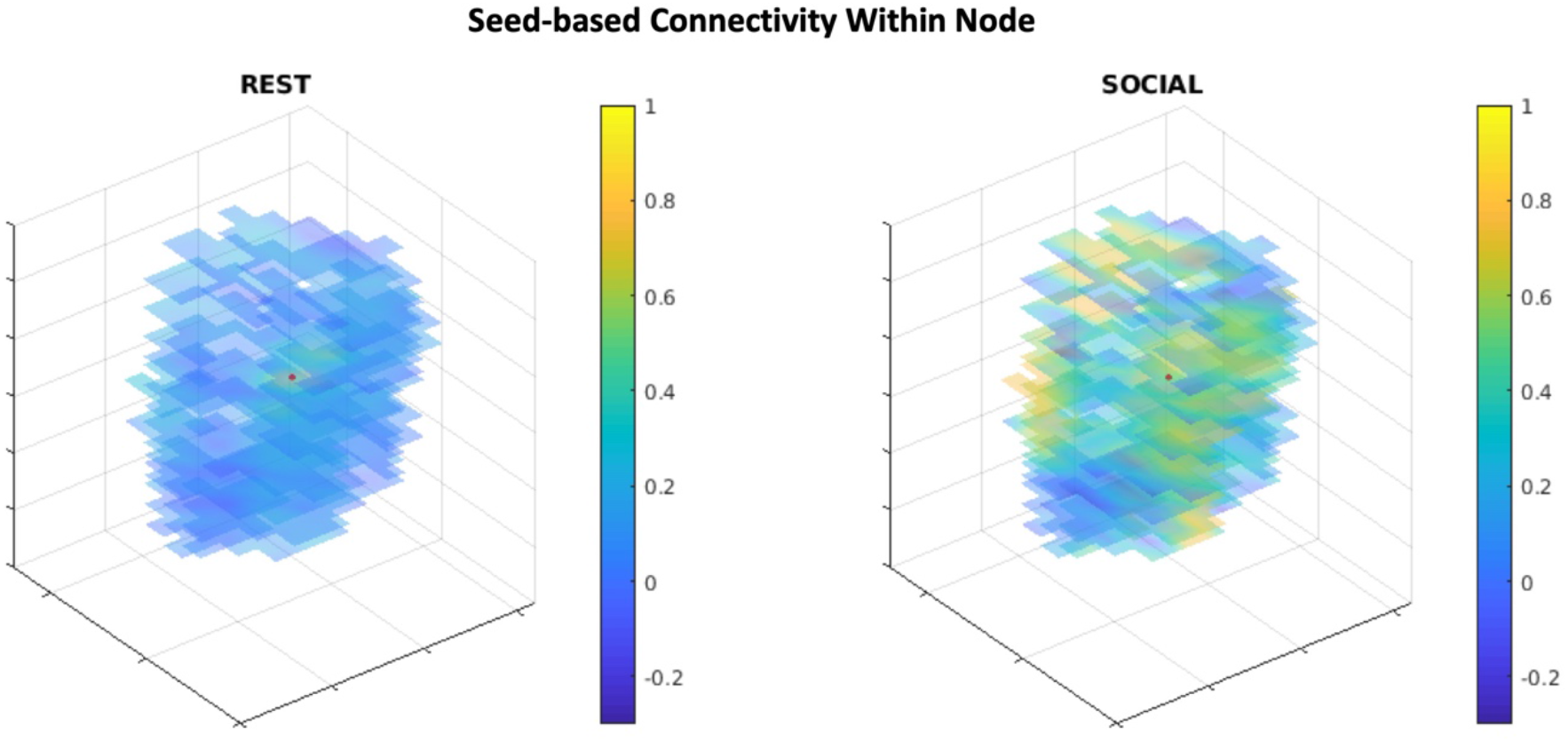
The between-voxel connectivity within Node 209 for subject 156536 under REST and SOCIAL. The seed voxel is marked by the red dot. The color of other voxels represents the functional connectivity between the voxel and the seed.

### 3.4 Subject groups defined by sex can be distinguished based on homogeneity vectors

Figure 4 demonstrates that using homogeneity as features, the gradient boosting classification model can classify sex at an accuracy much greater than chance. The classification accuracies, between 71% and 81%, are similar for all three atlases and all the tasks. The accuracies obtained from the random permutation test were approximately 50% as expected.

**Figure 4.**
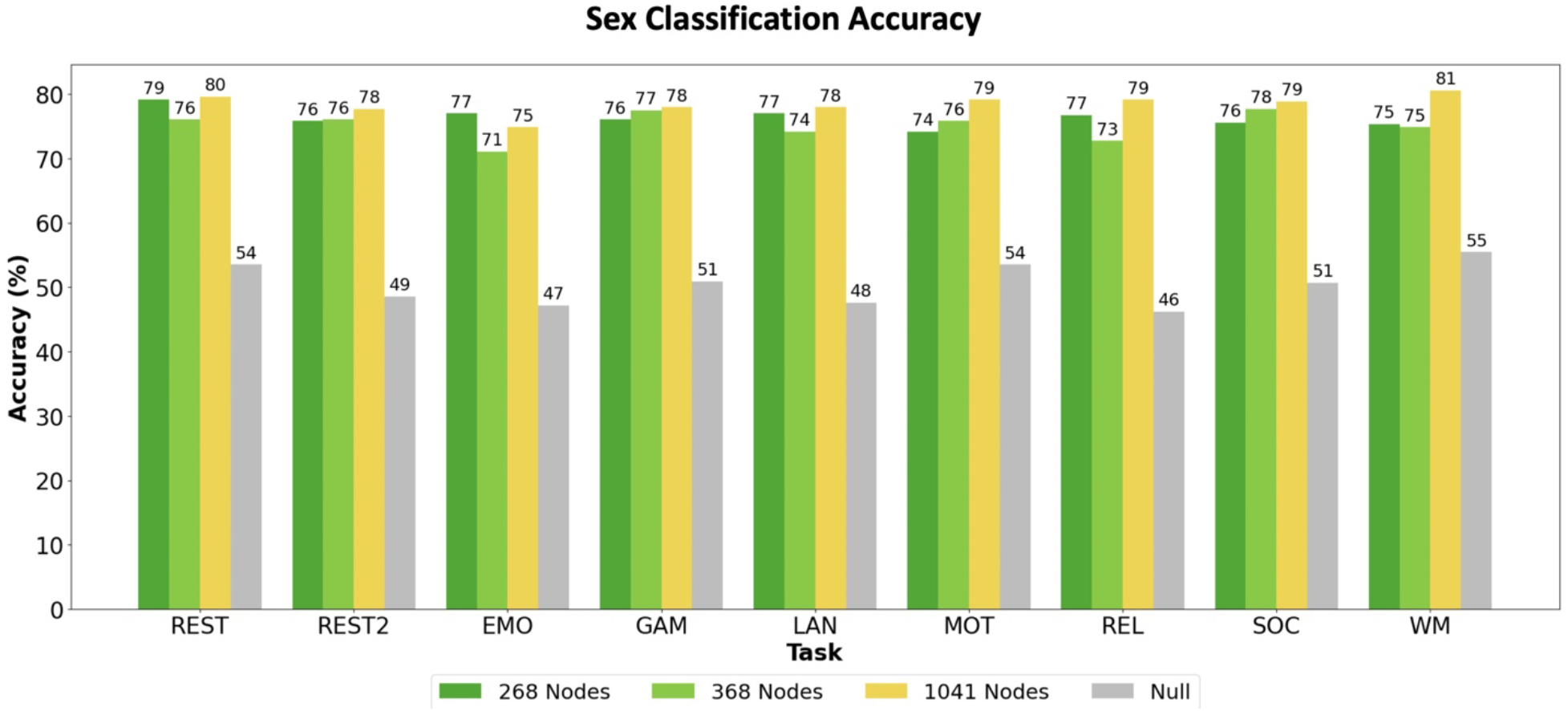
The sex classification accuracies for each task. Separate sex classification models were built and tested for each task with scans from 211 male and 211 female subjects. The accuracies for three different parcellations, Shen 268, Shen 368, and Yeo 1041 are reported in different colors. The gray bars represent the accuracies obtained from the permutation test where the sex labels are shuffled.

### 3.5 Subjects can be identified based on homogeneity vectors

We performed subject identification by evaluating the similarity of homogeneity vectors and quantified the accuracies of identification via these homogeneity vectors. (Figure 5a) Three sets of homogeneity vectors based on different atlases were used respectively. The identification accuracy should always be 1 if the database homogeneity vectors and target homogeneity vectors are from the same task, so the diagonals of all three accuracy matrices are 1. For Shen 268 (left), using GAMBLING as the database and WM as the target yields the highest accuracy, followed by REST2-REST, REST-REST2, and RELATIONAL-WM. The accuracy ranges widely from 0.16 to 0.82. For Shen 368 (middle), the overall accuracy increases while the relative accuracy, i.e. which pairs yield higher accuracies, stays similar. The lowest accuracy is 0.38 and the highest accuracy reaches 0.96. For Yeo 1041 (right), the accuracies for all task pairs are high, between 0.89 and 1. There is a clear increase in subject identification accuracy as the number of nodes in the atlas, i.e. the number of elements in the homogeneity vector increases.

**Figure 5.**
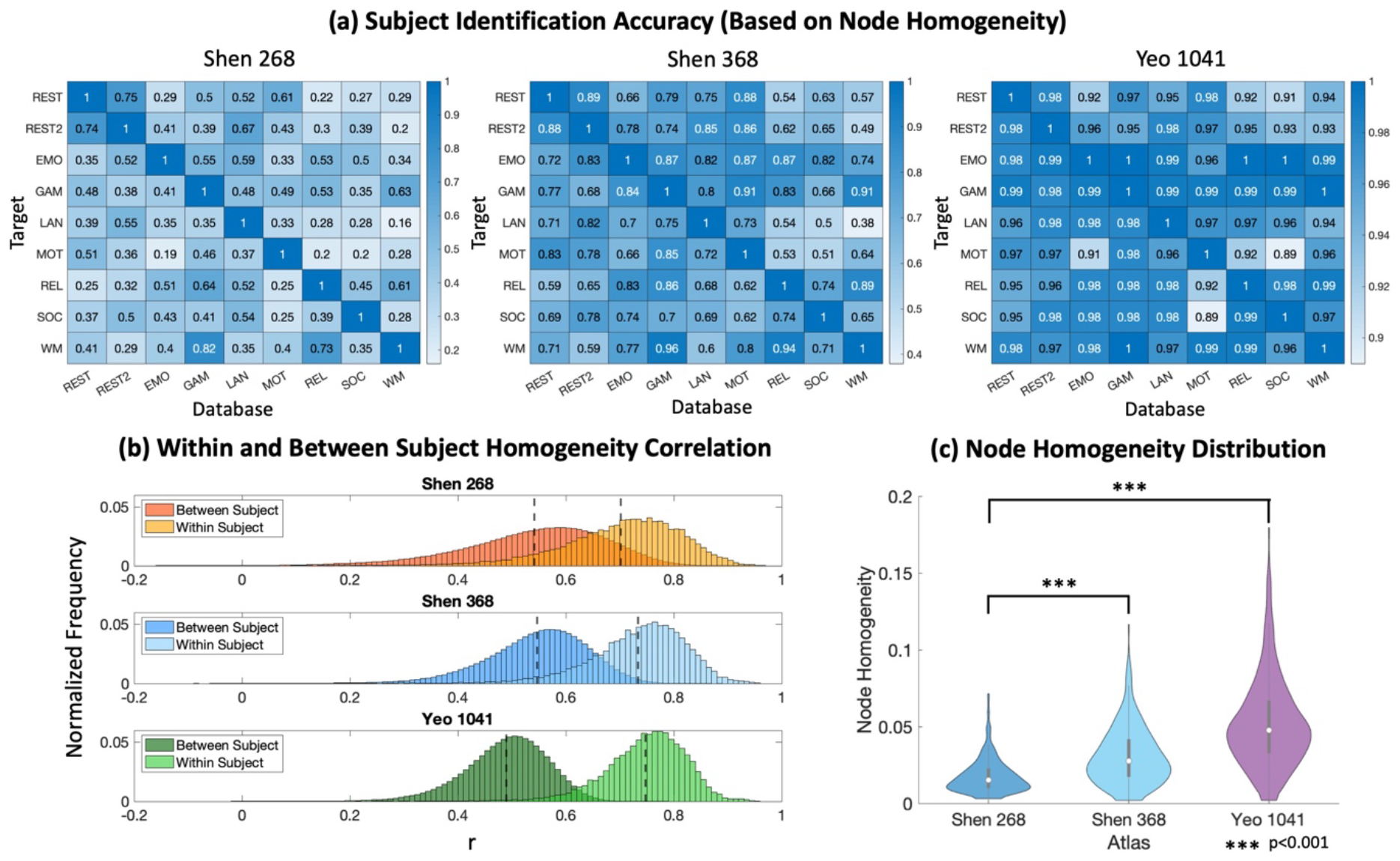
(a) The subject identification accuracies using homogeneity vectors based on Shen 268, Shen 368, and Yeo 1041. The x axis shows homogeneity vectors from which task are used as the database (Task X shown in Figure 1d) and the y axis shows homogeneity vectors from which task are used as the targets. The identification accuracy should always be 1 if the database homogeneity vectors and target homogeneity vectors are from the same task, so the diagonals of all three accuracy matrices are 1. Darker color represents higher accuracy. (b) The distribution of correlation coefficients between homogeneity vectors from the same subject and different subjects. (c) The distribution of node homogeneity for different atlases. The homogeneity for each node is averaged across subjects and states. The distributions are compared with F test.

We further explored the reason for this increase in accuracy. Figure 5b shows the distribution of correlation coefficients between homogeneity vectors from the same subject and different subjects. As the number of elements in homogeneity vectors increases, the standard deviations of the correlation coefficient between homogeneity vectors from the same subject and different subjects both decrease. Meanwhile, the correlation coefficient for within-subject homogeneity vector pairs increases as the number of nodes increases from 268 to 368 and 368 to 1041 (p<0.001). The correlation coefficient for between-subject homogeneity vector pairs is slightly higher for Shen 368 than for Shen 268 (p<0.001), but is much lower for Yeo 1041 than both Shen 268 and Shen 368 (p<0.001). In general, it is easier to distinguish between within-subject homogeneity pairs and between-subject homogeneity pairs as the number of nodes increases, which is consistent with the increase in identification accuracy in 4a.

The node homogeneity distributions for different atlases shown in Figure 5c indicate that as the number of nodes increases and node size decreases, the node homogeneity increases. The homogeneities are averaged across subjects and states. However, there are still nodes with very low homogeneity scores when the number of nodes in the applied atlas is relatively high. In all cases the nodes are far from being functionally homogeneous, i.e. having homogeneities of 1, yet they are not simply noisy but have reliable patterns of connectivity within them that allow both brain-state (task) classification and subject identification.

### 3.6 Homogeneity vectors based on state-specific and individualized parcellations are still informative

Individualized parcellations obtained with the exemplar-based individualized parcellation method attempt to account for between-voxel connectivity changes within nodes across subjects and states by allowing node reconfiguration. Similarly, state-specific parcellations capture node reconfiguration only across states driven by changes in voxel-to-voxel connectivity. If the flexible parcellations were ideal they would yield completely homogeneous nodes, of the form that did not change with changes in brain state or group. To test this theory, we computed the homogeneity vectors based on state-specific and individualized parcellations and tested whether they still have predictive power for task-induced brain states and subject identity.

Figure 6a shows the task classification accuracy using homogeneity vectors based on the fixed atlas (Shen 268), state-specific parcellations, and individualized parcellations. The accuracies for state-specific parcellations are about the same level as those for the fixed atlas. The accuracies decrease when homogeneity vectors based on individualized parcellations are used, especially for the tasks whose accuracies are the lowest among all the tasks (except for REST and REST2), GAMBLING, and RELATIONAL. Nonetheless, all the accuracies remain much higher than chance.

**Figure 6.**
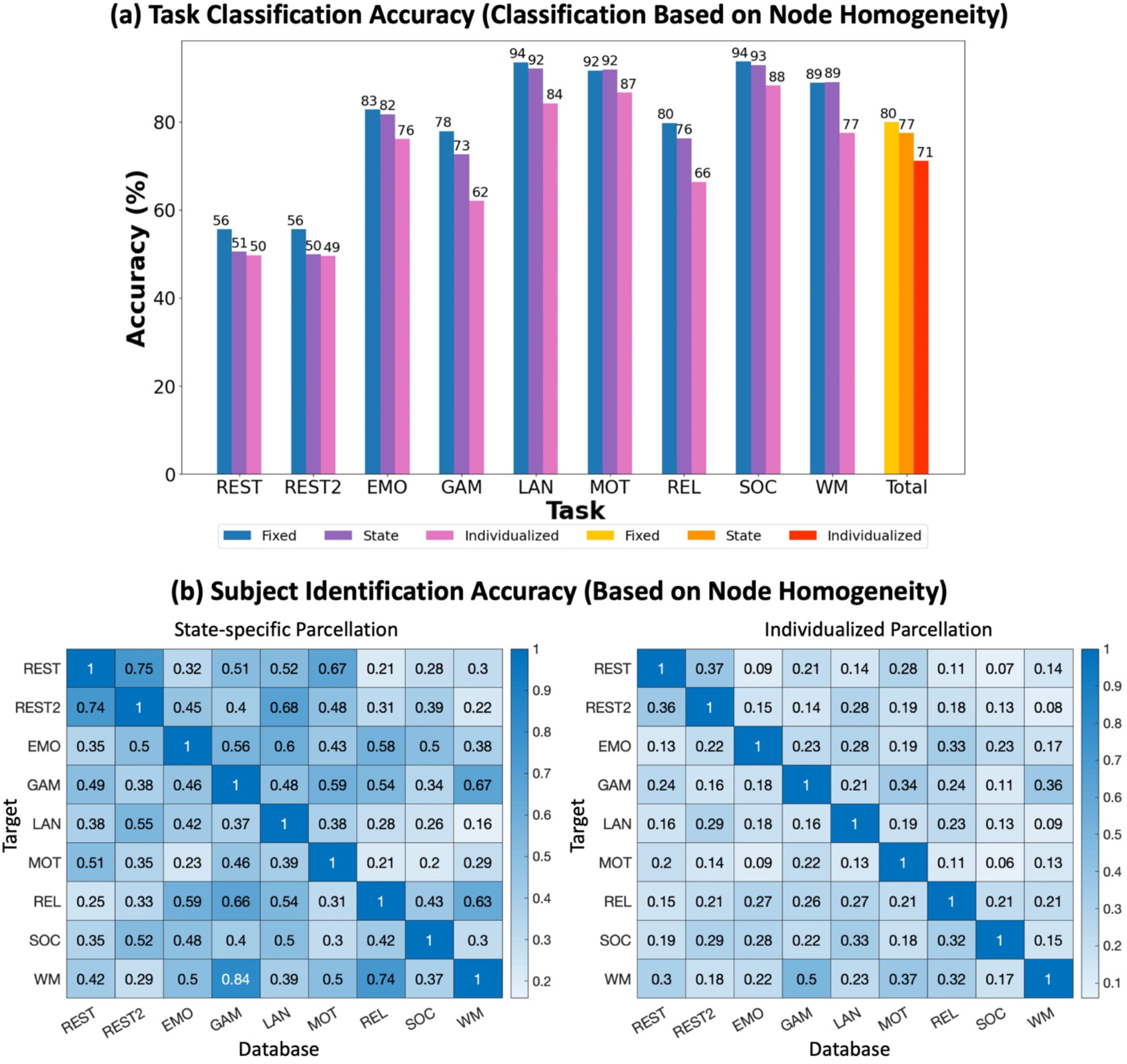
(a) The multiclass task classification accuracy using homogeneity vectors based on the fixed atlas (Shen 268), state-specific parcellations, and individualized parcellations. (b) The subject identification accuracies using homogeneity vectors based on state-specific parcellations and individualized parcellations.

The subject identification accuracies using homogeneity vectors based on state-specific parcellations (Figure 6b left) are very close to the accuracies for fixed atlas (Figure 5a left). The accuracies for individualized parcellations (Figure 6b right) are much lower than accuracies for the other two conditions.

We then compared the mean homogeneities across subjects and states based on different parcellation methods. Most of the data points representing homogeneities based on state-specific parcellations vs. fixed atlas (green dots in Figure 7a left) sit around the unity line indicating that the differences between homogeneities based on fixed and state-specific parcellations are small (mean = 6.5*10^−4^). The data points representing homogeneities based on individualized parcellations vs. fixed atlas (red dots) are mostly above the unity line indicating that homogeneities based on individualized parcellations are generally higher than those based on fixed parcellations. The mean difference is 0.0124. The data points with darker colors represent homogeneities of nodes that are more important in task classification. The results suggest that the important nodes tend to have higher homogeneities, which is consistent with the observation in Figure 3c. The right panel shows the distributions of homogeneities based on the three different parcellation methods. Node homogeneities based on state-specific parcellations and individualized parcellations are both higher than homogeneities based on fixed parcellations (p<0.001) although the differences between fixed homogeneities and state-specific homogeneities are small.

**Figure 7.**
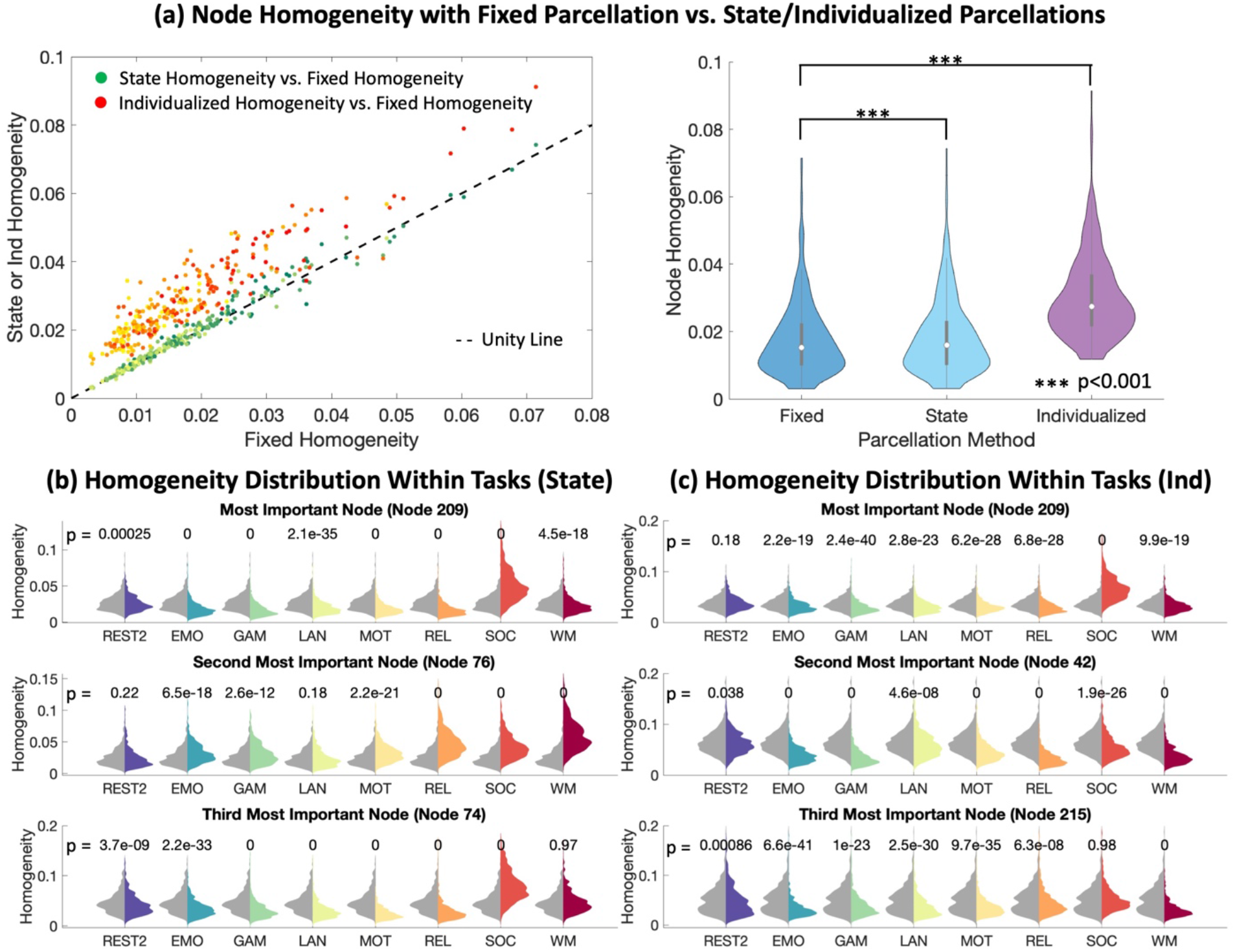
(a) Left panel: The green distribution shows the mean node homogeneity across subjects and states based on state-specific parcellations vs. fixed atlas (Shen 268). The orange distribution shows the mean node homogeneity based on individualized parcellations vs. fixed atlas. The dashed line is the unity line on which the homogeneities based on state-specific parcellations or individualized parcellations are the same as the homogeneities based on the fixed atlas. Right panel: The distribution of node homogeneity for fixed atlas, state-specific parcellations, and individualized parcellations. The homogeneity for each node is averaged across subjects and states. The distributions are compared with paired t-tests. (b) The homogeneity distributions based on state-specific parcellations for the three most important nodes (for task classification) are shown for all task conditions. The gray distributions represent the homogeneity distributions for REST for 493 subjects and are provided for reference, while the distributions in the other colors represent the homogeneities under tasks labeled on the x axis. Paired t-tests were used to compare the homogeneity distributions under REST and tasks. The p values are reported (above each pair of distribution). (c) The distributions of homogeneities based on individualized parcellations for the three most important nodes under different tasks.

Figure 7b and 7c show that the distributions of the three node homogeneities most important for task-classification are shown for both state-specific parcellations and individualized parcellations are significantly different during REST and other tasks. Many of the distributions are visually distinct. This could be the reason for the relatively high task classification accuracies shown in Figure 6a.

### 3.7 Homogeneity vectors based on intersect parcellation can be used for brain states decoding

Intercept parcellation contains only the voxels that are assigned to the same node across all nine state-specific parcellations. In Figure 8a, the REST parcellation is used as the reference and the ratio between the number of voxels in intersect parcellation and the number of voxels in REST parcellation is reported for each node. The ratio ranges from 0.43 to 0.96 with a mean of 0.80. This indicates that the majority of the voxels are assigned to the same node across tasks.

**Figure 8.**
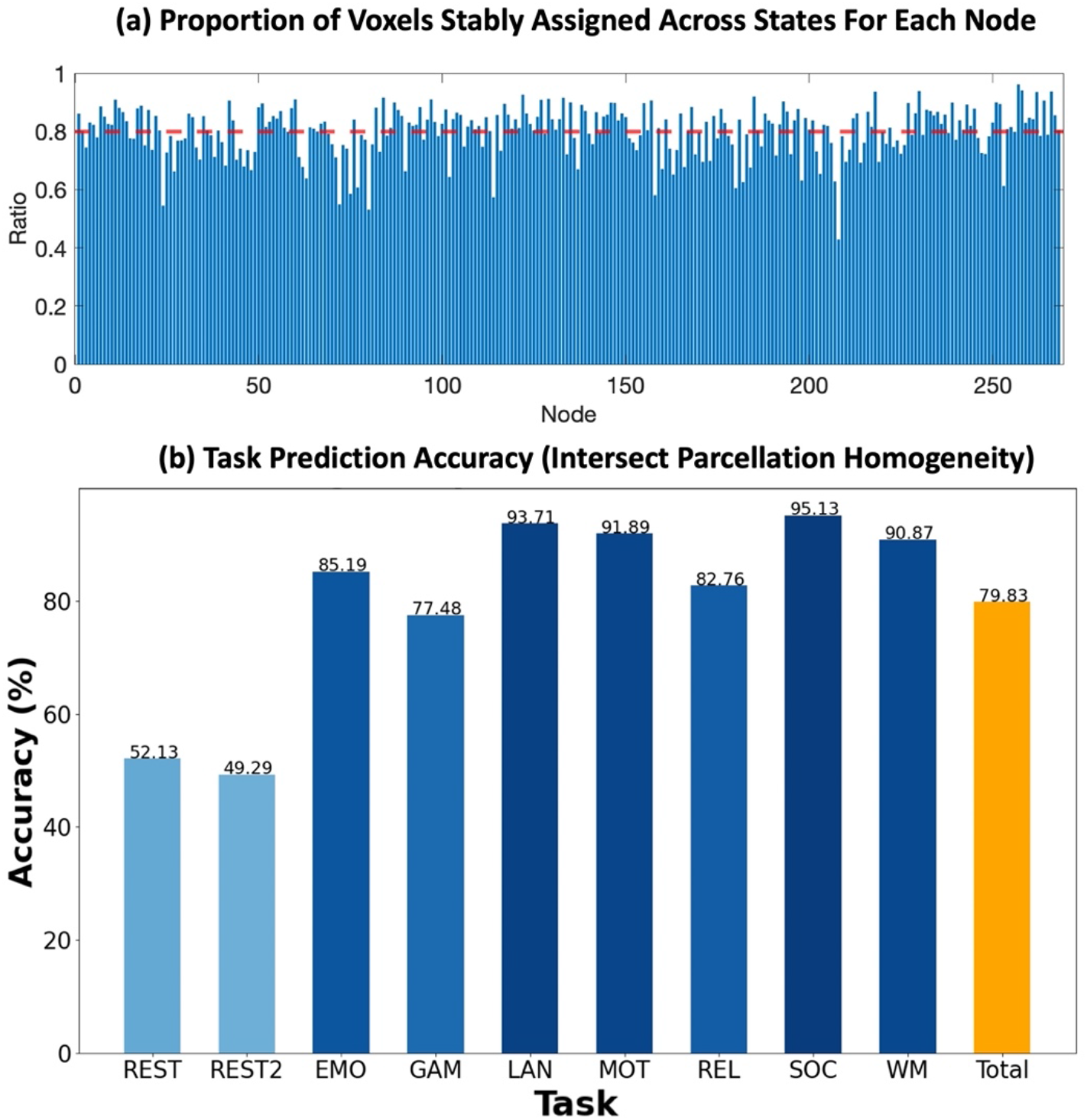
(a) The proportion of voxels stably assigned to the same node across all states among all voxels for each node in REST state-specific parcellation, i.e. the ratio between node size in the intersect parcellation and REST state-specific parcellation. The red dashed line shows the mean ratio across all the nodes. (b) The multiclass task classification accuracies for each task and the total accuracy using homogeneity vectors based on the intersect parcellation.

To test whether the between-voxel connectivity of such a group of stably assigned voxels still has predictive power for tasks, we performed the same task classification based on homogeneity as described above. In this case, each node contains fewer voxels. Figure 8b shows that the task classification accuracies remain high using homogeneity vectors based on the intersection of parcellations, comparable with the accuracies based on fixed atlas.

## Discussion

### 4.1 Predictive power of within-node functional connectivity

Node homogeneity, which is the mean of the within-node functional connectivity, is a coarse metric. Yet, here we have shown that it can be used to classify tasks, distinguish groups defined by sex, and identify subjects with high accuracy. These findings demonstrate that the functional connectivity within-nodes changes systematically with task-conditions and subject groups, and also provides additional information unique to each subject. Within-node connectivity, co-varying with task or group is typically not measured nor controlled for in connectivity analyses. Detecting edge level connectivity changes in situations where the source of change may be within-node connectivity reconfigurations could provide an incomplete picture of the connectivity landscape, when contrasting groups or tasks.

There are several plausible reasons for the systematic changes in node homogeneity. The nodes in the most commonly used atlases may contain multiple distinctive, functional subunits that have specialized tunings, that vary with task. If the brain has the order of 16 billion neurons, then a 400-node atlas contains of the order of 40 million neurons. It is not a stretch to imagine that these neurons are not all tuned to the same inputs and outputs. Yet using a fixed atlas in functional connectivity work makes exactly that assumption.

The standard approach of averaging the time series within the node not only obscures this potentially interesting within-node dynamic organization but can also distort the connectivity between nodes. One of the fixed atlases we applied, Yeo 1041 contains more nodes than most atlases. The homogeneities based on Yeo 1041 are higher than those based on atlases with fewer nodes. Yet, the task classification accuracy using homogeneity vectors based on this atlas is comparable to the accuracies based on the other two atlases and much higher than chance. This suggests that a 1041 node atlas still contains functional subunits. Given that functional organization in the brain is flexible^33^, and that functional units can be overlapping^39^, it is not clear that a fixed atlas at any scale would overcome this problem, except at the limit of course, treating each voxel as a node yields only 1 within-node time-course.

Across all 3 atlases tested, the Shen 268, Shen 368, and Yeo 1041, the subject identification accuracy increases as the number of nodes increases likely because the number of features in the vector increases. In the limit of completely homogeneous nodes, there should be no information in the homogeneity vector, and subject identification accuracy should be at chance even if the atlas resolution is increased further.

### 4.2 Homogeneities based on flexible parcellations and the intersect parcellation

With state-specific parcellations, nodes are allowed to reconfigure, in a data driven manner, across different task-induced brain states. Individualized parcellations provide even more flexibility because each parcellation is derived for each subject and each state. However, the results show that the homogeneity vectors based on state-specific parcellations and individualized parcellations can still be used to predict tasks indicating that the variance in within-node connectivity across states is not fully accounted for by these flexible parcellations (Figure 6a). Subject identification accuracies are almost the same with fixed atlas and state-specific parcellations, suggesting that although the parcellation applied for each state is unique, the homogeneity patterns for different subjects are preserved. The subject identification accuracies based on individualized parcellations are lower than those based on the other two parcellation methods (Figure 6b) possibly because allowing the parcellations to vary across subjects breaks the subject-specific patterns in homogeneity vectors. Individualized parcellations can also be noisier because they are based on a relatively small amount of data.

The information provided by within-node connectivity remains even with the state-specific and individualized parcellations possibly because of the restricted extent to which these parcellations are allowed to deviate from the initial fixed atlas. The exemplar-based individualized parcellation method preserves the node correspondence between individualized parcellations, which means the number of nodes is always held constant. The exemplar for each node is identified within the node in the initial fixed atlas and is usually near the geometric center of the node. In addition, the voxel-to-parcel assignment has a spatial contiguity constraint. Due to these restrictions, most of the voxels (∼80%) around the geometric center of the nodes are consistently assigned to the same node across all state-specific parcellations. However, as shown in Figure 3, the changes in connectivity within a node across states take place throughout the node. Such changes are not fully accounted for with the individualized parcellation method applied here.

To investigate performance of a parcellation that was consistent across tasks, we generated an intersect-parcellation, whose nodes contained only voxels that were consistently assigned to the same node across all state-specific parcellations. Each node in the intersect parcellation contains 43% to 96% of the voxels in the original REST state-specific parcellation, with a mean of 80% (Figure 8b). The connectivity within-nodes in this intersect-parcellation however, still varies systematically across tasks, suggesting that an alternative approach is needed to fully account for these within-node connectivity changes.

### 4.3 Individualized parcellation methods account for inter-state and inter-subject differences

Studies have shown that there can be differences in the spatial configurations of nodes across subjects ^44^ and individualized parcellation methods have been developed to attempt to account for these differences^45, 46^. Methods have also been developed to compensate for both spatial differences between subjects as well as task-induced brain states^33, 34^. Individualized parcellations have been shown to have higher functional homogeneity than fixed atlases^33, 45, 46^. The subject-specific parcellations are relatively stable for the same subject and distinctive across different subjects^46^. Subtle differences in parcellations have been related to cognitive measures^45^, and a metric extracted from subject-specific state-specific parcellations, the node size vector, has been shown to be useful in predicting task performance^33^.

The evidence strongly supports that functional spatial configurations vary systematically across subjects and states, yet the field has been slow to adapt to this functional flexibility. While the individualized atlas approaches attempt to accommodate functional reconfiguration, they do not achieve uniform state-independent homogeneity and sub-specialization within the individualized nodes remains. There is no consensus on which individualized parcellation method is the optimal and open questions remain as to the ideal number of nodes, and whether or not these should be contiguous. The change in homogeneity vector with task could be added as a constraint to a parcellation algorithm, where parcel size is decreased until the within-node connectivity between tasks stabilizes. Here we tested the implicit assumption of the fixed atlas approach that the node can be treated as a homogeneous unit, and we demonstrate that significant changes in connectivity occur within-nodes in a reliable manner. The work here does not demonstrate, nor argue, that one atlas is better than another. But it does indicate that the use of a fixed atlas when studying changes in connectivity across brain states or groups, overlooks significant within node changes in connectivity. Such changes occur even with atlases of up to 1000 nodes, which is a higher resolution than found in most studies.

### 4.4 Implications

Almost all studies to date have used atlases with hard boundaries and employed the same atlas across all subjects and, in studies where task data is included, all task-conditions. The underlying assumption has been that the node acts as a cohesive unit and that within-node connectivity is high, constant, and therefore not of interest. The homogeneity measure used in this work is only a coarse metric to measure the extent to which voxels within a node share the same temporal dynamics across scan conditions. This coarse measure however contains sufficient information to allow task classification (gradient boosting classifier), classification of subjects by sex, and subject identification (correlation-based fingerprinting), and yet these are by no means the optimal models to extract information on within node connectivity changes. The results suggest that a considerable amount of information on the connectivity topology is missed when only edge changes are considered. A danger in not considering within node connectivity changes is that such changes may be incorrectly ascribed to external edge changes leading to incorrect interpretation of the connectivity changes observed.

Although state-specific parcellations have previously been shown to reflect dynamic brain functional organization ^33^, the results here show that they do not fully account for changes in within-node connectivity to yield completely homogeneous parcellations that do not vary systematically with task or group. However, there are several potential directions for future works.

First, increasing the number of nodes in the atlas does lead to higher within-node functional similarity. Since one possible reason for the systematic changes in node homogeneity is the merging of different functional subunits, further dividing the nodes to separate these subunits is a potential solution. It remains an open question whether there exist spatially contiguous units that are consistently functionally synchronized under all states at the current scale of neuroimaging. It is shown in this work that parcellations with as many as 1041 nodes are not fine enough to ensure stable within-node connectivity. Previous work ^33^ has revealed consistent and predictive reconfiguration for parcellations containing even 5000 nodes. Thus, if there exists a fixed atlas that meets the previously described assumption, it is likely to require even more than 5000 nodes.

Allowing more flexible parcellations is another potential way to better account for within-node connectivity variance. Current approaches to flexible parcellation hold the number of nodes fixed and limit the amount of reconfiguration in order to maintain node correspondence. It is currently unknown however, if the number, size and location of functional units varies substantially across subjects and brain states. However, to compare connectomes across individuals and tasks requires node correspondence across all these conditions. Thus, it can be challenging to find the balance between these competing goals for parcellation.

For some analyses, metrics derived from within-node connectivity, such as homogeneity, could be included as separate factors. For example, homogeneity vectors can be included as an additional set of input variables when building models to predict behavior from connectivity. It is not straightforward however to incorporate metrics such as homogeneity in direct contrasts of connectivity matrices or networks. Even if this is not achievable, at a minimum, the variance of within-node connectivity should not be correlated with the variable of interest. For example, if one were to investigate differences in node-level functional connectivity between task-induced states, within-node connectivity should not vary systematically with task-induced states. This study, however, demonstrates that using current methods it does co-vary with the variables of interest.

Finally, if we consider an atlas only as a tool for dimension reduction, then simply performing the analyses on voxel-level connectivity rather than node-level connectivity mostly avoids the challenge of choosing the appropriate parcellation. Previous studies have investigated voxel-level connectivity analysis^47, 48^, and some metrics based on voxel-level connectivity have been proposed to prevent the high computational expense of full voxel-level connectivity matrices^49^.

## Conclusions

In this work, we demonstrate that connectivity between voxels within-nodes varies systematically and predictively across subjects and task-induced brain states. Almost all functional connectivity studies to date have used fixed parcellations across a range of conditions or groups, focusing only on changes in node-to-node connectivity while ignoring within-node connectivity changes, which are shown here to be informative. Such changes in within-node connectivity may be misinterpreted as edge changes leading to erroneous conclusions. The results are shown to replicate across three different fixed atlases covering resolutions from 268 to over 1000 nodes. Several potential strategies can be adopted to account for the within-node connectivity changes and make use of this additional information when studying connectivity changes between groups and/or different task conditions and brain states.

## Acknowledgment

Support from NIH MH121095 is gratefully acknowledged

## Methods

### 2.1 Data

A subset of the Human Connectome Project (HCP) S900 release was used. Only subjects who had voxel-level fMRI data for all nine functional sessions (two resting-states and seven tasks) with left-right phase-encoding were included. To alleviate artifacts caused by head motion, subjects with mean frame-to-frame displacement > 0.1 mm or maximum frame-to-frame displacement > 0.15 mm were excluded. The resulting dataset contains 493 subjects (266 females, age = 22-36+). The preprocessing procedures were the same as described in ^33^. We applied the HCP minimal preprocessing pipeline ^50^ which includes artifact removal, motion correction, and registration to MNI space. All further preprocessing steps used BioimageSuite ^51^, including regressing 24 motion parameters, regressing the mean time courses of the white matter and cerebrospinal fluid, and the global signal, removing the linear trend and low pass filtering.

### 2.2 Brain atlases and parcellations

Three different functional atlases were used to investigate whether the results depend on atlas choice. Shen 268 is a 268-parcel atlas determined with a spectral clustering algorithm on resting-state data of a healthy population ^25^. Shen 368 is a finer atlas obtained by integrating the parcellation of cortex from ^25^, subcortex from the anatomical Yale Brodmann Atlas ^52^, and cerebellum from Yeo et al. ^26^. Yeo 1041 was obtained by integrating the subcortical and cerebellum parcellation of the 368-parcel parcellation with the 1000-parcel cortex parcellation from Yeo et al. ^26^.

The exemplar-based individualized parcellation method previously proposed in ^33^ was used to generate the individualized parcellations for each scan. The algorithm has three steps: 1) Registration to a group-level parcellation. Here we used the Shen 268 atlas as the initial group-level parcellation. 2) Identification of an exemplar from each node by maximizing a submodular function. 3) Assignment of each voxel to the functionally closest exemplar while maintaining spatial contiguity to an exemplar.

The state-specific parcellations were determined by taking the majority vote over the individualized parcellations for all the subjects during the same task. For example, a voxel in the working memory parcellation is assigned to the node to which the voxel is most frequently assigned across all 493 subject-specific working memory individualized parcellations.

Based on the nine state-specific parcellations, we obtained an intersect parcellation which only contains voxels that are assigned to the same node across all state-specific parcellations. Ambiguous voxels which are assigned to different nodes under different states were not included in this parcellation leaving uncovered gaps between nodes. The intersect parcellation covers approximately 79% of the voxels covered in the resting-state state-specific parcellation.

### 2.3 Homogeneity

We used node homogeneity to summarize between-voxel connectivity within-nodes and investigated how it changes across states and subjects. The homogeneity of each node is defined as the mean pair-wise correlation between voxel-level time series within the node. There is a homogeneity vector associated with each scan which contains the homogeneities of all the nodes defined by the atlas applied. The dimension of a homogeneity vector depends on the resolution of the atlas. (Figure 9a) Six sets of homogeneity vectors were computed based on different atlases (Shen 268, Shen 368, Yeo 1041, state-specific parcellations, individualized parcellations, and intersect parcellation).

**Figure 9.**
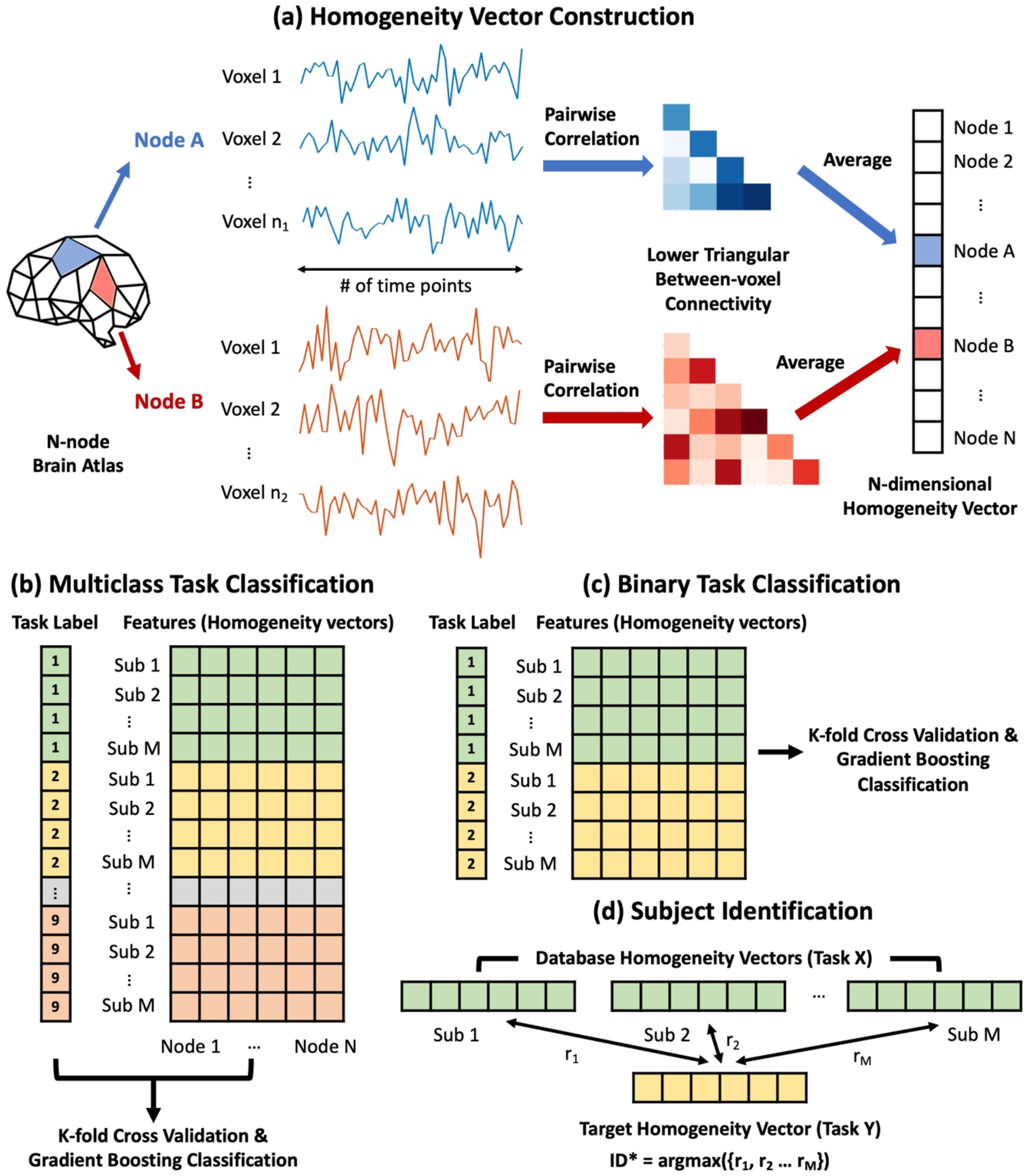
(a) The homogeneity of each node is defined as the mean pair-wise correlation between voxel-level time series within the node. The within-node between-voxel connectivity matrices can have different sizes because the nodes can contain different numbers of voxels. There is a homogeneity vector associated with each scan which contains the homogeneities of all the nodes defined by the atlas applied. The number of elements in the homogeneity vector is the same as the number of nodes in the atlas applied. (b) Each scan is considered as one entry. The scans are labeled 1 to 9 depending on the task performed during the scans. All the entries (493 subjects * 9 sessions) are divided into k folds. At each iteration, a gradient-boosting classifier was trained with k-1 folds and then used to predict the task labels for the remaining fold. (c) Binary task classification is almost the same as multiclass task classification in (b) except that only scans from two different tasks are being distinguished each time. (d) To identify the subject using a homogeneity vector, we first built a database containing homogeneity vectors for all subjects from task X. Then, the similarity between the target homogeneity vector from task Y and every vector in the database was evaluated. The predicted ID of the target vector would be the same as its most similar vector in the database. Here the similarity between vectors was evaluated by Pearson correlation.

### 2.4 Task classification using homogeneity vectors as features

We built and tested a gradient-boosting classifier (GBM; with 300 estimators and learning rate = 0.1) model that predicts task-induced brain states using homogeneity vectors as features. With the k-fold cross-validation approach (k = 10 in this study), all the samples/scans (493 subjects * 9 sessions) are divided into k folds. At each iteration, the model was trained with k-1 folds and then used to predict the task labels for the remaining fold. (Figure 9b) After N iterations, the predicted labels were compared to the true labels. The predictive power was evaluated by the mean predictive accuracy for each of the tasks, i.e. the proportion of correctly labels scans among all the scans whose true labels are task X, and the total accuracy. The accuracies range between 0% and 100% and higher accuracy indicates stronger predictive power. The importance of the homogeneity of each node for task classification is evaluated by the impurity-based feature importance extracted from the gradient-boosting classifier.

A permutation test was performed by randomly shuffling the task labels of the scans before building the gradient-boosting classifier and evaluating the predictive power in the same way as described above. Homogeneity vectors based on Shen 268 were used in the permutation test.

We also performed binary task classifications where each sample is labeled one of the two tasks rather than nine tasks. For each task pair, 986 samples (493 subjects * 2 sessions) were used for the same cross-validation and model building procedure. (Figure 9c) A total of 36 (9*(9-1)/2) accuracies for all task pairs were then computed and reported.

### 2.5 Sex classification using homogeneity vectors as features

The same gradient-boosting classifier model introduced in 2.4 was trained to classify subject sex based on homogeneity vectors. 211 male and 211 female subjects with matched age distributions were included. For each task, 422 scans were used for cross-validation and model training.

### 2.6 Statistical test comparing homogeneity distributions

For each node, the distributions of homogeneity across subjects under different states (tasks) are compared using paired t-test.

The distributions of mean homogeneity across subjects and states for different fixed atlases (Shen 268, Shen 368, and Yeo 1041) are compared using F test because the numbers of nodes are different for different parcellations. The distributions of mean homogeneity for fixed atlas (Shen 268), state-specific parcellation, and individualized parcellation are compared using paired t-test which is possible because this exemplar based parcellation maintains node correspondence across different parcellations.

### 2.7 Subject Identification using homogeneity vectors

To identify the subject using a homogeneity vector, we first built a database containing homogeneity vectors for all subjects from task X. Then, the similarity between the target homogeneity vector from task Y and every vector in the database was evaluated. The predicted ID of the target vector would be the same as its most similar vector in the database. Here the similarity between vectors was evaluated by Pearson correlation. (Figure 9d) A higher correlation coefficient r indicates higher similarity. The associated ID of each target vector was predicted independently, so multiple target vectors can be assigned the same ID although the true ID of every target vector from task Y is unique. A similar subject “fingerprinting” was performed in ^8^ except that between-node connectivity was used as the feature in this previous study.

The task pairs used as the database and target are referred to as task X-task Y in Results and Discussion, e.g. REST-EMOTION means homogeneity vectors from REST are used as the database and homogeneity vectors from EMOTION are the targets whose ID are being predicted.

### 2.8 Between-voxel connectivity visualization

To better investigate the topography of voxel-to-voxel connectivity within a node, we randomly selected a seed voxel near the center of a node, for those nodes whose homogeneity was highly predictive for tasks. The seed-voxel to other voxels connectivity within the same node was calculated and visualized for different tasks in order to visualize within-node connectivity topology.

## References

1. Sporns O, Tononi G, Kötter R. The human connectome: A structural description of the human brain. PLoS Computational Biology 1, 0245–0251 (2005).

2. Biswal B, Yetkin FZ, Haughton VM, Hyde JS. Functional connectivity in the motor cortex of resting human brain using echo-planar MRI. Magnetic resonance in medicine : official journal of the Society of Magnetic Resonance in Medicine / Society of Magnetic Resonance in Medicine 34, 537–541 (1995).

3. Rubinov M, Sporns O. Complex network measures of brain connectivity: uses and interpretations. NeuroImage 52, 1059–1069 (2010).

4. Satterthwaite TD, et al. Linked Sex Differences in Cognition and Functional Connectivity in Youth. Cereb Cortex 25, 2383–2394 (2015).

5. Zhang C, Cahill ND, Arbabshirani MR, White T, Baum SA, Michael AM. Sex and age effects of functional connectivity in early adulthood. Brain connectivity 6, 700–713 (2016).

6. Geerligs L, Renken RJ, Saliasi E, Maurits NM, Lorist MM. A Brain-Wide Study of Age-Related Changes in Functional Connectivity. Cereb Cortex 25, 1987–1999 (2015).

7. Santarnecchi E, Emmendorfer A, Tadayon S, Rossi S, Rossi A, Pascual-Leone A. Network connectivity correlates of variability in fluid intelligence performance. Intelligence 65, 35–47 (2017).

8. Finn ES, et al. Functional connectome fingerprinting: identifying individuals using patterns of brain connectivity. Nature neuroscience 18, 1664–1671 (2015).

9. Rosenberg MD, Finn ES, Scheinost D, Constable RT, Chun MM. Characterizing Attention with Predictive Network Models. Trends in cognitive sciences 21, 290–302 (2017).

10. Rosenberg MD, et al. A neuromarker of sustained attention from whole-brain functional connectivity. Nature neuroscience 19, 165–171 (2016).

11. Rosenberg MD, et al. Functional connectivity predicts changes in attention observed across minutes, days, and months. Proceedings of the National Academy of Sciences of the United States of America 117, 3797–3807 (2020).

12. Tian L, Wang J, Yan C, He Y. Hemisphere-and gender-related differences in small-world brain networks: a resting-state functional MRI study. Neuroimage 54, 191–202 (2011).

13. Meunier D, Achard S, Morcom A, Bullmore E. Age-related changes in modular organization of human brain functional networks. Neuroimage 44, 715–723 (2009).

14. Rudie JD, et al. Altered functional and structural brain network organization in autism. NeuroImage: clinical 2, 79–94 (2013).

15. Cohen JR, D’Esposito M. The segregation and integration of distinct brain networks and their relationship to cognition. Journal of Neuroscience 36, 12083–12094 (2016).

16. Shine JM, et al. The dynamics of functional brain networks: integrated network states during cognitive task performance. Neuron 92, 544–554 (2016).

17. Eickhoff SB, Constable RT, Yeo BT. Topographic organization of the cerebral cortex and brain cartography. Neuroimage 170, 332–347 (2018).

18. Brodmann K. Vergleichende Lokalisationslehre der Großhirnrinde: in ihren Prinzipien dargestellt auf Grund des Zellenbaues. Barth JA (1909).

19. Blumensath T, et al. Spatially constrained hierarchical parcellation of the brain with resting-state fMRI. Neuroimage 76, 313–324 (2013).

20. Smith SM, et al. Functional connectomics from resting-state fMRI. Trends in cognitive sciences 17, 666–682 (2013).

21. Fan L, et al. The human brainnetome atlas: a new brain atlas based on connectional architecture. Cerebral cortex 26, 3508–3526 (2016).

22. Glasser MF, et al. A multi-modal parcellation of human cerebral cortex. Nature 536, 171–178 (2016).

23. Gordon EM, Laumann TO, Adeyemo B, Huckins JF, Kelley WM, Petersen SE. Generation and evaluation of a cortical area parcellation from resting-state correlations. Cerebral cortex 26, 288–303 (2016).

24. Power JD, et al. Functional network organization of the human brain. Neuron 72, 665–678 (2011).

25. Shen X, Tokoglu F, Papademetris X, Constable RT. Groupwise whole-brain parcellation from resting-state fMRI data for network node identification. Neuroimage 82, 403–415 (2013).

26. Yeo BT, et al. The organization of the human cerebral cortex estimated by intrinsic functional connectivity. Journal of neurophysiology, (2011).

27. Rolls ET, Huang C-C, Lin C-P, Feng J, Joliot M. Automated anatomical labelling atlas 3. Neuroimage 206, 116189 (2020).

28. Caviness Jr VS, Meyer J, Makris N, Kennedy DN. MRI-based topographic parcellation of human neocortex: an anatomically specified method with estimate of reliability. Journal of cognitive neuroscience 8, 566–587 (1996).

29. Messé A. Parcellation influence on the connectivity-based structure–function relationship in the human brain. Human brain mapping 41, 1167–1180 (2020).

30. Park B, Ko JH, Lee JD, Park H-J. Evaluation of node-inhomogeneity effects on the functional brain network properties using an anatomy-constrained hierarchical brain parcellation. PloS one 8, e74935 (2013).

31. Qi S, Meesters S, Nicolay K, ter Haar Romeny BM, Ossenblok P. The influence of construction methodology on structural brain network measures: A review. Journal of neuroscience methods 253, 170–182 (2015).

32. Lord A, et al. Brain parcellation choice affects disease-related topology differences increasingly from global to local network levels. Psychiatry Research: Neuroimaging 249, 12–19 (2016).

33. Salehi M, Greene AS, Karbasi A, Shen X, Scheinost D, Constable RT. There is no single functional atlas even for a single individual: Functional parcel definitions change with task. NeuroImage 208, 116366 (2020).

34. Boukhdhir A, Zhang Y, Mignotte M, Bellec P. Unraveling reproducible dynamic states of individual brain functional parcellation. Network Neuroscience 5, 28–55 (2021).

35. Calhoun VD, Kiehl KA, Pearlson GD. Modulation of temporally coherent brain networks estimated using ICA at rest and during cognitive tasks. Human brain mapping 29, 828–838 (2008).

36. Ma S, Calhoun VD, Phlypo R, Adali T. Dynamic changes of spatial functional network connectivity in healthy individuals and schizophrenia patients using independent vector analysis. NeuroImage 90, 196–206 (2014).

37. Iraji A, et al. The spatial chronnectome reveals a dynamic interplay between functional segregation and integration. Human brain mapping 40, 3058–3077 (2019).

38. McIntosh AR. Contexts and catalysts: a resolution of the localization and integration of function in the brain. Neuroinformatics 2, 175–182 (2004).

39. Yeo BT, et al. Functional Specialization and Flexibility in Human Association Cortex. Cerebral cortex (New York, NY : 1991) 25, 3654–3672 (2015).

40. Luo W, Greene AS, Constable RT. Within node connectivity changes, not simply edge changes, influence graph theory measures in functional connectivity studies of the brain. NeuroImage, 118332 (2021).

41. Bozek J, et al. Construction of a neonatal cortical surface atlas using Multimodal Surface Matching in the Developing Human Connectome Project. NeuroImage 179, 11–29 (2018).

42. Lawrence RM, et al. Standardizing human brain parcellations. Scientific Data 8, 1–9 (2021).

43. Schaefer A, et al. Local-global parcellation of the human cerebral cortex from intrinsic functional connectivity MRI. Cerebral cortex 28, 3095–3114 (2018).

44. Bijsterbosch JD, et al. The relationship between spatial configuration and functional connectivity of brain regions. Elife 7, e32992 (2018).

45. Kong R, et al. Spatial topography of individual-specific cortical networks predicts human cognition, personality, and emotion. Cerebral cortex 29, 2533–2551 (2019).

46. Wang D, et al. Parcellating cortical functional networks in individuals. Nature Neuroscience 18, 1853–1860 (2015).

47. Hayasaka S, Laurienti PJ. Comparison of characteristics between region-and voxel-based network analyses in resting-state fMRI data. Neuroimage 50, 499–508 (2010).

48. Wang Y, Cohen JD, Li K, Turk-Browne NB. Full correlation matrix analysis (FCMA): An unbiased method for task-related functional connectivity. J Neurosci Methods 251, 108–119 (2015).

49. Scheinost D, et al. The intrinsic connectivity distribution: a novel contrast measure reflecting voxel level functional connectivity. Neuroimage 62, 1510–1519 (2012).

50. Glasser MF, et al. The minimal preprocessing pipelines for the Human Connectome Project. Neuroimage 80, 105–124 (2013).

51. Joshi A, et al. Unified framework for development, deployment and robust testing of neuroimaging algorithms. Neuroinformatics 9, 69–84 (2011).

52. Lacadie CM, Fulbright RK, Rajeevan N, Constable RT, Papademetris X. More accurate Talairach coordinates for neuroimaging using non-linear registration. Neuroimage 42, 717–725 (2008).

